# The chloroplast-localised transporter RETICULATA controls palisade mesophyll cell division and expansion

**DOI:** 10.64898/2026.11.03.759069

**Authors:** Edwyn A. Locke Beviá, Emilio Aldorino, James M. Fitzsimons, Jean C. Dillon, Chris D. Whitewoods

## Abstract

Plant leaves contain extensive intercellular air spaces, which contribute to photosynthetic capacity and water use efficiency in different environments. Despite this importance, the developmental mechanisms that control leaf air space patterning are poorly understood. For over 20 years it has been known that plants with disrupted chloroplast biogenesis and metabolism have increased intercellular air spaces, but the underlying molecular mechanism is unknown. Here, we characterise leaf air space morphogenesis in mutants of the chloroplast amino acid transporter *RETICULATA* (*RE*), which has enlarged air spaces. We show that the *reticulata-6* (*re-6*) loss of function mutant exhibits increased air space expansion specifically in the palisade, and not the spongy, mesophyll. We also show that *RE* does not regulate air space patterning via programmed cell death, via a change in stomatal density or a change in cell identity of the mesophyll tissue. Instead, we quantify mesophyll cell morphology and division rates to show that RE acts by promoting early palisade cell division and suppressing later cell expansion. Overall, we show that RE specifically regulated palisade mesophyll morphogenesis and air space patterning and hypothesise that chloroplasts may act to regulate air space patterning in the palisade mesophyll in response to light.

## INTRODUCTION

Leaves are full of air. In terms of volume, intercellular air spaces (IAS) in the leaves of vascular plants range from 3% (Slaton & Smith, 2002) to 73% (Earles *et al*., 2018). The development of IAS was a key innovation during the transition from aquatic to terrestrial habitats because the resistance for CO_2_ diffusion is ∼10,000-fold lower in air than in water (Earles *et al*., 2018), which meant plants could profit from an increased photosynthetic rate. However, despite the central role of IAS in maximising leaf photosynthetic capacity (Lehmeier *et al*., 2017; Mathers *et al*., 2018), how IAS patterning unfolds during leaf development remains largely unresolved.

Many leaves contain two distinct photosynthetic cell layers: the palisade and the spongy mesophyll (Pyke *et al*., 1991; Bar & Ori, 2014). The palisade layer is located beneath the adaxial epidermis and is characterised by having small air spaces and tightly packed cylindrical shaped cells, optimised for efficient light harvesting (Terashima & Inoue, 1984). In contrast, the spongy mesophyll, which is between the palisade mesophyll and the abaxial epidermis, has larger air spaces and irregularly shaped lobed cells that enhance gas exchange and light scattering (DeLucia *et al*., 1996; Smith *et al*., 1997). This coordinated tissue architecture integrates light capture with internal CO_2_ diffusion.

Throughout the mesophyll, IAS are formed by cell separation at multicellular junctions (Avery Jr, 1933; Jeffree *et al*., 1986; Fox *et al*., 2018). After initial formation, IAS expansion is controlled separately in the palisade and spongy. In the palisade, further cell separation (schizogeny) is regulated in part by growth of the constraining epidermis (Fitzsimons *et al*., 2026), whereas in the spongy, air spaces enlarge by preferential expansion of the air space adjacent mesophyll cell wall (expansigeny) (Pyke *et al*., 1991; Jarvis *et al*., 2003; Zhang *et al*., 2021).

Signals originating from chloroplasts (Li *et al*., 1995; Hricová *et al*., 2006; Pérez-Pérez *et al*., 2013; reviewed in Lundquist *et al*., 2014) and stomata (Masle *et al*., 2005; Dow *et al*., 2017; Lundgren *et al*., 2019; reviewed in Baillie & Fleming, 2020) are also known to alter IAS patterning, but the molecular and developmental mechanisms by which they work are unknown.

Mutants in chloroplast biogenesis and metabolism have enlarged IAS with irregularly shaped palisade mesophyll cells (Li *et al*., 1995; Chatterjee *et al*., 1996; Keddie *et al*., 1996; Voll *et al*., 2003). The best studied of these is the mutant *reticulata* (*re*) (Barth & Conklin, 2003; González-Bayon *et al*., 2006; Pérez-Pérez *et al*., 2013; Kuhnert *et al*., 2025), which was identified by George Rédei in 1964 (Rédei & Hironyo, 1964). This mutant is named for its distinctive leaf colouration of marked dark green veins on a pale green lamina (Fig. 1). *RE* is a nuclear gene that encodes a basic amino acid transporter localised to the inner plastid envelope, and homologs of this protein are solely found in plastid containing organisms (Kuhnert *et al*., 2025). *RE* is mainly expressed in the proximal region of young leaves, whereas in older leaves expression is observed in the bundle sheath and surrounding cells, and in the hydathodes (Pérez-Pérez *et al*., 2013). *re* mutants show no detectable differences in chloroplast number or morphology relative to wild-type (Barth & Conklin, 2003; González-Bayon *et al*., 2006), meaning that its effects on leaf structure are likely through changes in metabolites or molecular signals rather than through lack of chloroplast function.

**Fig. 1.**
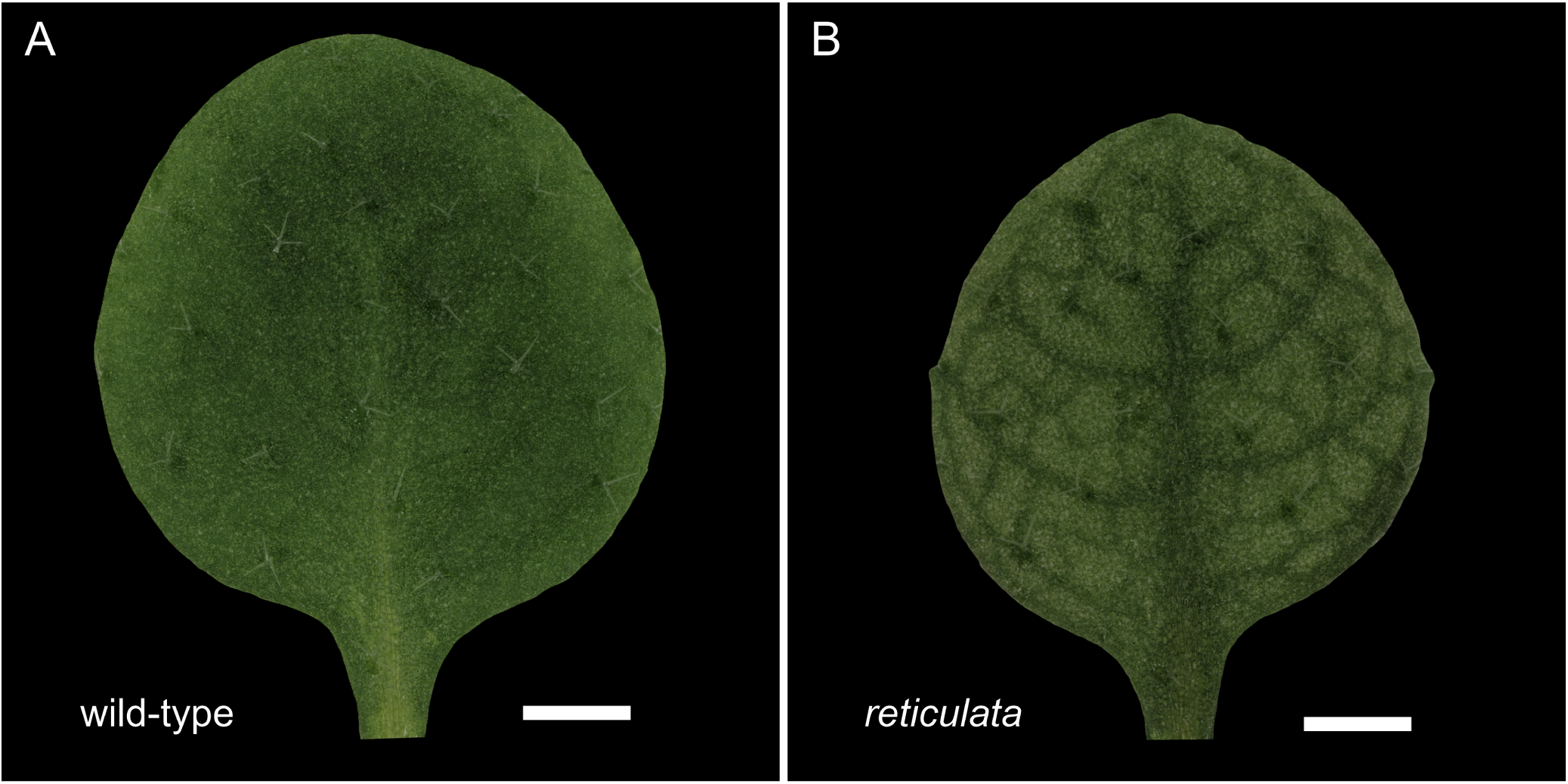
The *reticulata* mutant is characterised by a dark green venation on a pale green lamina. (A) First leaf of a 23 days after stratification (DAS) *A. thaliana* wild-type plant. (B) First leaf of a 23 DAS *A. thaliana re-6* mutant plant. Scale bars: 1 mm.

Although the mechanism for IAS enlargement in the *re* mutant is not known, it has been proposed that it is caused by mesophyll cell death due to increased reactive oxygen species (ROS) in mature leaves (Pérez-Pérez *et al*., 2013). In this view, the enhanced air spaces are a pathological outcome of chloroplast dysfunction, and RE (and chloroplasts) may have no physiological role in IAS patterning in wild-type. However, cell death has not been observed in *re* mutant palisade cells, and ROS has been linked to altered cell expansion during leaf development (Schmidt *et al*., 2016). Therefore, we hypothesise that enlarged IAS in *re* are not due to cell death but are instead an outcome of either 1– altered palisade division and expansion or 2– a change in adaxial mesophyll cell identity from palisade to spongy. Either of these possibilities would suggest that RE actively regulates IAS patterning.

Here we investigate how *RE* controls air space development. We use confocal microscopy to visualise IAS development in *re-6* mutant leaves throughout development, quantify cell death using fluorescent dyes and use cell type-specific markers to observe any changes in palisade and spongy mesophyll identity. We show that the *re-6* mutant’s enlarged air spaces are specific to palisade tissue and form gradually throughout development as a result of reduced palisade cell division, rather than through cell death or a switch in cell identity. This suggests that RE is required for mesophyll morphogenesis and IAS patterning.

## RESULTS

### The *re-6* mutation specifically disrupts IAS and cell morphology in the palisade mesophyll

To compare IAS expansion during leaf development in both *A. thaliana* wild-type and *re-6* mutant plants, we quantified porosity (percentage of air space) in the first rosette leaf pair across different timepoints, from initial air space formation to leaf maturity. We performed our analysis on the first two true leaves because they are initiated almost simultaneously at opposite positions of the SAM (Van Lijsebettens & Clarke, 1998), and share the reticulate phenotype with older leaves. To quantify porosity we used two different dyes: propidium iodide or Nile Red. For leaves sampled 8 days after stratification (DAS) we used Nile Red in Silicon oil, which enters through stomata and visualises IAS via green fluorescence (Kawase *et al*. (2015); Kawase *et al*. (2016); Fig. 2A,E). Prior to 8 DAS Nile Red does not stain IAS as stomata were not yet open. Instead, we used propidium iodide staining to highlight individual cells (grey), which contrasts with IAS (black) after a process of tissue fixation and clearing (see Materials and Methods and Fig. 2A,E).

**Fig. 2.**
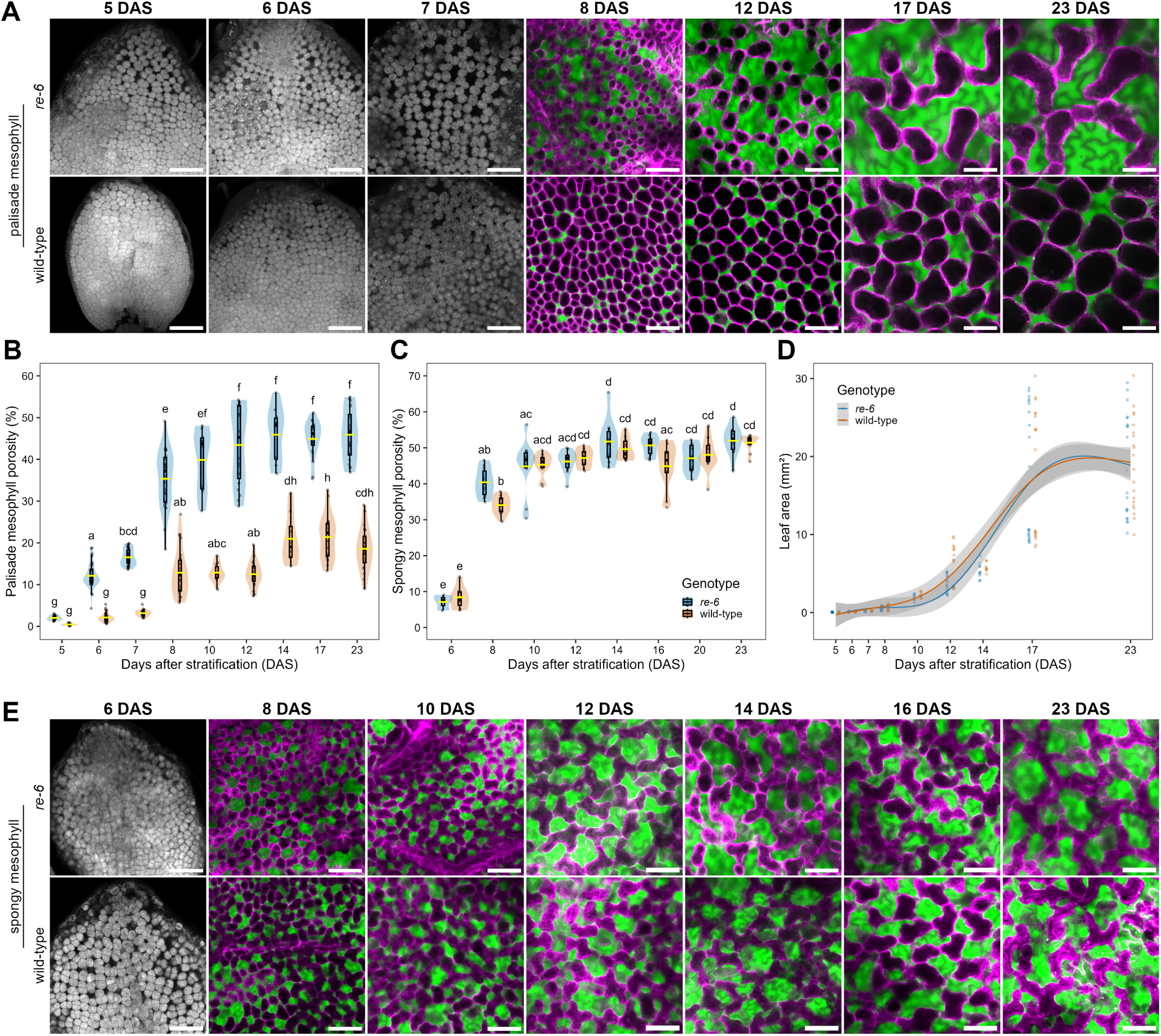
The *re-6* mutation disrupts palisade, but not spongy, mesophyll development. (A) Air spaces in the palisade mesophyll of *re-6* (top row) and wild-type (bottom row) visualised by propidium iodide (5–7 DAS) or Nile Red dye (8–23 DAS). (B,C) Palisade (B) and spongy (C) mesophyll tissue porosity (%) of wild-type and *re-6* first rosette leaf pair over 23 DAS. Significance letters above violins derive from a one-way ANOVA followed by Tukey’s HSD post-hoc test. Yellow horizontal lines on violins indicate mean values. (D) Progression of leaf area over time of *A. thaliana* first rosette leaf pair for wild-type and *re-6* with a generalised additive model (GAM) fit and 99% confidence interval. (E) Air spaces in the spongy mesophyll of *re-6* (top row) and wild-type (bottom row) visualised by propidium iodide (6 DAS) or Nile Red dye (8–23 DAS). With propidium iodide, air spaces are black and cells grey; with Nile Red air spaces are green and cell outlines magenta. Each point represents an individual leaf. Scale bars: 50 µm.

Since leaf development progresses basipetally (Pyke *et al*., 1991; Donnelly *et al*., 1999; Fox *et al*., 2018), we quantified porosity at the upper half of the leaf blade across the different timepoints (Fig. 2B,C). In the wild-type, and consistent with previous observations (Fitzsimons *et al*., 2026), IAS expansion in the palisade and spongy mesophyll followed distinct trajectories: IAS in the palisade increased slowly before reaching a plateau of ∼20% porosity at 14 DAS, whereas the spongy mesophyll expanded more rapidly and plateaued at ∼50% (Fig. 2B,C). The steepest phase of IAS expansion occurred between 7-8 DAS in the palisade (3 to 13%) and between 6-8 DAS in the spongy mesophyll (8 to 34%).

Although the growth of wild-type and *re-6* leaves was not significantly different (Fig. 2D), in *re-6*, the rate of IAS expansion in the palisade mesophyll was markedly accelerated (Fig. 2A,B), while the spongy mesophyll followed a wild-type-like pattern (Fig. 2C,E). Palisade porosity in *re-6* was only slightly elevated at 5 DAS (0.5% for wild-type and 2% for *re-6*), and from 6 DAS onward became significantly higher than in the wild-type, increasing from 12 to 45% by 14 DAS, compared with 2 to 20% in the wild-type. Thus, *re-6* palisade porosity is 2.3-fold higher than in the wild-type by the time it plateaus, despite both genotypes reaching a plateau at roughly the same time (14 DAS). Cell density and cell shape in the palisade mesophyll of *re-6* visually differ from wild-type, resembling the lobed morphology of spongy mesophyll cells (Fig. 2A,E). Together, these results suggest that the *re-6* mutation specifically disrupts palisade mesophyll development, affecting both IAS patterning and cell morphology.

### Altered adaxial-abaxial stomatal patterning does not account for increased air space in *re-6*

Previous work has shown that stomatal density is positively correlated with palisade mesophyll porosity (Dow *et al*., 2017; Lundgren *et al*., 2019). Therefore, to test whether the enlarged IAS in the *re-6* mutant palisade is a secondary effect driven by stomata, we quantified stomatal density (stomata/mm^2^) in both the adaxial and abaxial epidermis of the first rosette leaf pair of *A. thaliana* wild-type and *re-6* plants by using Nile Red staining to visualise stomata (Fig. 3A). We saw no significant difference in stomatal density between wild-type and *re-6* leaves other than at 10 DAS on the adaxial epidermis and at 8 DAS on the abaxial epidermis (Fig. 3B,C). In both cases, wild-type stomatal density was 16 and 19% lower, respectively, than in *re-6*. Given the transient nature of these differences during leaf development, these data suggest the specific increase in the palisade mesophyll porosity of *re-6* is not due to an indirect effect of altered stomatal patterning.

**Fig. 3.**
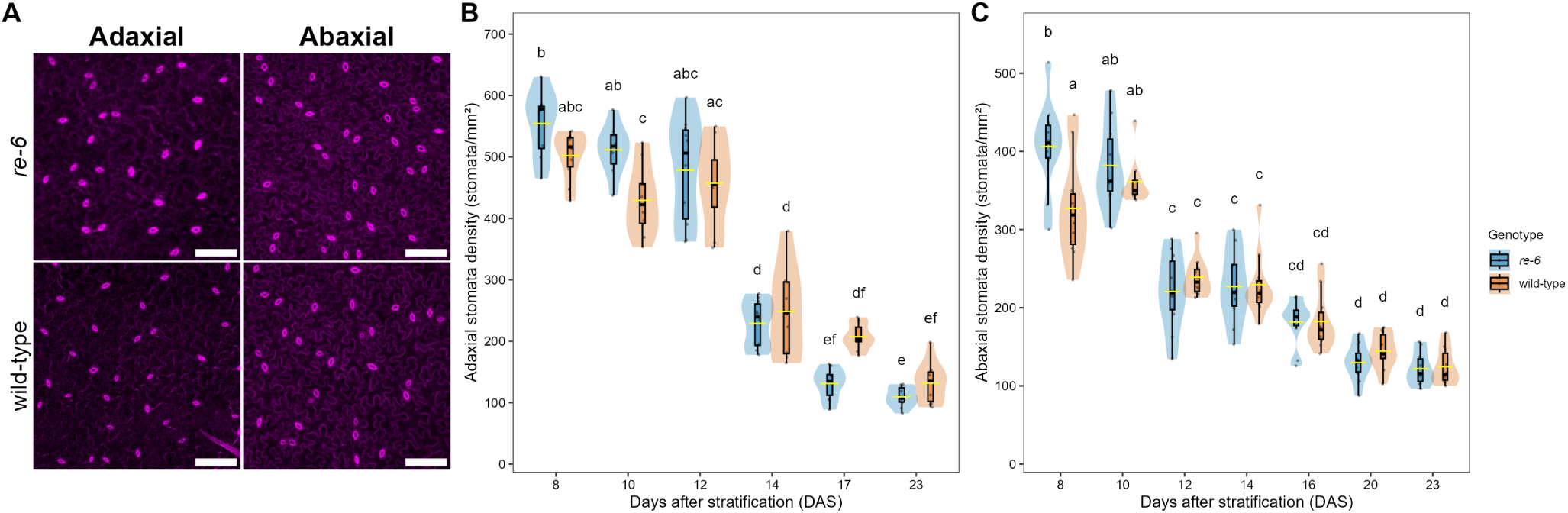
Stomatal density in *re-6* adaxial and abaxial epidermis is comparable to wild-type. (A) Representative adaxial and abaxial epidermis of 23 DAS *re-6* (top row) and wild-type (bottom row) visualised by Nile Red dye. Cell outlines magenta. (B,C) Adaxial (B) and abaxial (C) stomatal density (stomata/mm^2^) of wild-type and *re-6* first rosette leaf pair over 23 DAS. Significance letters above violins derive from a one-way ANOVA followed by Tukey’s HSD posthoc test. Yellow horizontal lines on violins indicate mean values. Each point represents an individual leaf. Scale bars: 100 µm.

### Palisade cell death does not contribute to IAS formation in *re-6*

Previous studies have proposed that the increased IAS volume observed in *re* mutants may result from mesophyll cell death associated with elevated ROS in mature leaves (Barth & Conklin, 2003; Overmyer *et al*., 2008; Pérez-Pérez *et al*., 2013). However, cell death has never been documented during *re* leaf development. Therefore, to determine if the mechanism for IAS enlargement in *re-6* is caused by palisade mesophyll cell death, we used Sytox Orange to stain dead cells in rosette leaf one of *A. thaliana* wild-type and *re-6* plants at 8 and 14 DAS (Fig. 4A). To verify we could reliably detect dead palisade cells with our assay, we performed targeted laser ablations of palisade cells followed by Sytox Orange staining. The ablated palisade cells were successfully stained, confirming the effectiveness of the staining method for detecting cell death in this tissue (Fig. S1). We detected no significant difference in the proportion of dead palisade cells between wild-type and *re-6* leaves at either timepoint. In wild-type leaves, dead cells represented 4% of palisade cells at 8 DAS and 0.4% at 14 DAS, whereas in *re-6*, dead cells accounted for 2% of palisade cells at 8 DAS, and no dead palisade cells were detected at 14 DAS (Fig. 4B). Together, these results indicate that the specific increase in palisade mesophyll IAS in *re-6* is not caused by cell death. Instead, they suggest RE may play a physiological role in IAS patterning in wild-type leaves.

**Fig. 4.**
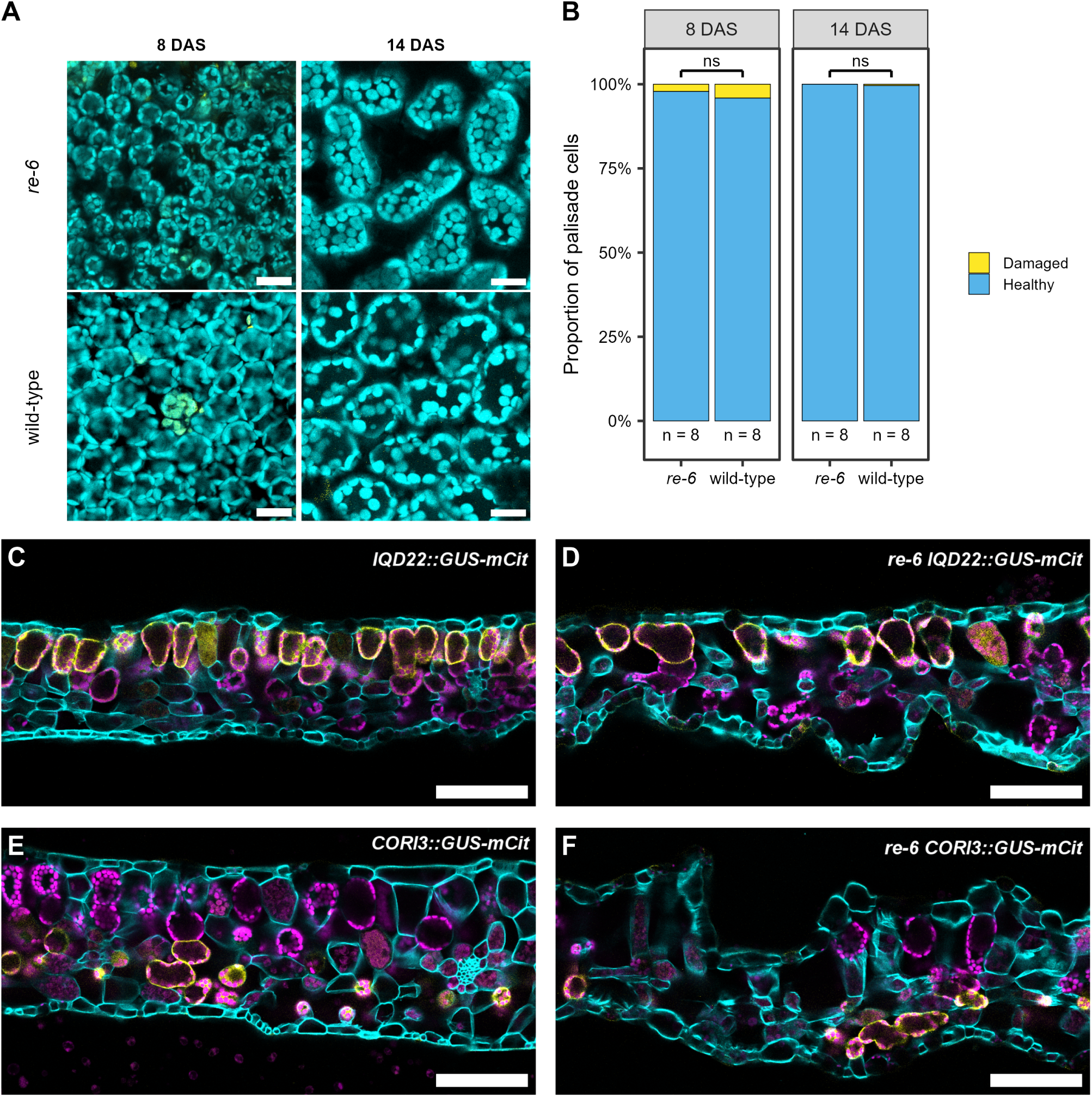
Increased IAS in the *re-6* palisade is not caused by cell death or a switch in cell identity. (A) Representative images of palisade mesophyll of rosette leaf one stained with Sytox Orange at 8 and 14 DAS (left and right columns, respectively). *re-6* (top row) and wild-type (bottom row). Chloroplast autofluorescence is visualised in cyan and Sytox Orange in yellow. Scale bars: 20 µm. (B) Proportion of damaged and healthy palisade mesophyll cells of wild-type and *re-6* rosette leaf one at 8 and 14 DAS. Statistical significance was calculated with Welch’s two-sample t-test (unequal variances): *P<0.05, **P<0.01, ***P<0.001, ns = not significantly different. (C-F) Representative images of transverse sections of leaf one from wild-type *IQD22::GUS-mCit* (C), *re-6 IQD22::GUS-mCit* (D), wild-type *CORI3::GUS-mCit* (E) and *re-6 CORI3::GUS-mCit* (F). Calcofluor white is visualised in cyan, mCitrine in yellow and chloroplast autofluorescence in magenta. Scale bars: 100 µm.

### Increased air space in *re-6* is not caused by a switch in cell identity

As palisade mesophyll cell morphology in *re-6* at the end of leaf expansion resembles the lobed spongy mesophyll cells (Fig. 2A, E), we hypothesised that the enlarged IAS in *re-6* might result from a switch in mesophyll cell identity from palisade to spongy. To determine if this is the case, we examined palisade and spongy cell identity in wild-type and *re-6* leaves at 14 DAS using the cell type-specific reporters *IQD22::GUS-mCit* and *CORI3::GUS-mCit*, which mark palisade (Fig. 4C) and spongy (Fig. 4E) mesophyll cells, respectively (Procko *et al*., 2022). *IQD22* expression in the *re-6* mutant remained detectable in palisade cells (Fig. 4D), indicating retention of palisade cell identity. Likewise, *CORI3* expression remained confined to spongy mesophyll cells in *re-6*, whereas palisade cells lacked detectable expression (Fig. 4F) despite their spongy-like morphology. These data suggest that palisade mesophyll cell identity in *re-6* has not switched to spongy mesophyll. Therefore, the increased palisade mesophyll IAS observed in *re-6* cannot be explained as a result of altered mesophyll cell identity.

### The *re-6* mutation reduces palisade cell proliferation rate and alters cell morphology

Since *re-6* appeared to have altered palisade mesophyll cell density and morphology (Fig. 2A), we quantified these traits throughout development of the first rosette leaf pair of *A. thaliana* wild-type and *re-6* plants to determine if the enlarged IAS in *re-6* palisade could be explained by reduced cell density in this tissue and to assess to what extent the *re-6* mutation affects palisade cell morphology. As above, quantifications were carried out in the upper half of the leaf blade (Fig. 5B-F) using CellposeSAM v4.0.4 cellular segmentation software (Pachitariu *et al*., 2025) on images of palisade mesophyll stained with Nile Red (as in Fig. 2A). An example of the cellular segmentation of the palisade mesophyll for a 23 DAS wild-type leaf can be seen in Fig. 5A.

**Fig. 5.**
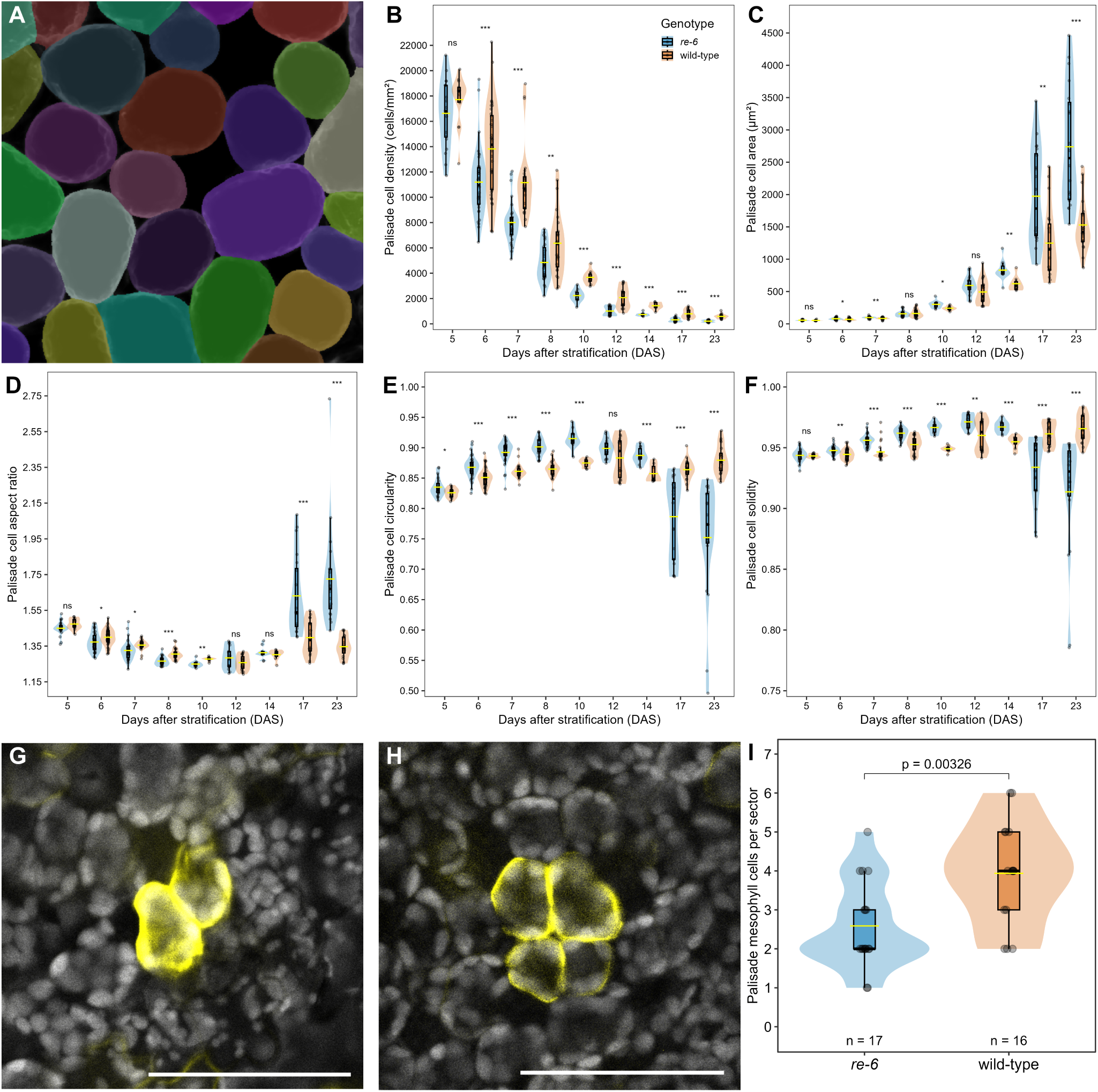
Palisade mesophyll cells of *re-6* divide more slowly and are larger than wild-type. (A) Snapshot of the palisade mesophyll of a 23 DAS wild-type leaf segmented with CellposeSAM v4.0.4. (B–F) Palisade cell (B) density (cells/mm^2^), (C) area (µm^2^), (D) aspect ratio, (E) circularity and (F) solidity of wild-type and *re-6* first rosette leaf pair over 23 DAS. Each data point represents the mean value for a leaf. (G, H) Wild-type (G) and *re-6* (H) palisade mesophyll clonal sectors expressing YFP in a 7 DAS leaf 3 days after heat-shock induction. Chlorophyll autofluorescence is visualised in grey, YFP in yellow. (I) Quantification of cells in clonal sectors expressing YFP in palisade mesophyll of 7 DAS leaves induced at 4 DAS. Data represent individual sectors. Statistical significance was calculated with Welch’s two-sample t-test (unequal variances): *P<0.05, **P<0.01, ***P<0.001, ns = not significantly different. Yellow horizontal lines on violins indicate mean values. Scale bars: 50 µm.

In the *re-6* mutant, the previously observed accelerated rate of IAS expansion in the palisade mesophyll (Fig. 2A,B) was matched by a proportional and gradual decrease in cell density throughout leaf development relative to the wild-type (Fig. 5B). As with palisade porosity, differences in cell density arose at 6 DAS. From this timepoint onward, *re-6* consistently exhibited lower cell density, reaching a 2.8-fold reduction compared with the wild-type by 23 DAS (Fig. 5B). Given the similar leaf area, these findings suggest that the enlarged IAS phenotype in *re-6* palisade mesophyll could be explained by a reduced proliferation rate of the palisade cells. To test whether the palisade mesophyll cell division rate was decreased in *re-6*, we performed clonal analysis by using heat shock-inducible CRE-mediated lox site recombination to induce YFP expression (Fig. 5G, H) (Kuchen *et al*., 2012; Lee *et al*., 2019; Fitzsimons *et al*., 2026). We induced YFP expression in single cells and counted the number of cells in each clonal sector 3 days later as a proxy for cell division rate. These data show that cell number per sector was significantly different between wild-type (3.94 cells/sector) and *re-6* (2.59 cells/sector; Fig. 5I), the latter having on average 1.35 less cells per sector compared to the wild-type. This is equivalent to 0.63 divisions per day in wild type and 0.42 divisions per day in *re-6*, indicating that the proliferation rate of palisade cells in *re-6* is around 30% lower than that of the wild-type in the 4 to 7 DAS period of leaf development. Notably, reduction of palisade cell division rate in *re-6* occurs right before the steepest phase of IAS expansion (7-8 DAS), when IAS go from 17 to 35% (Fig. 2B). Thus, it seems likely that the enlarged IAS phenotype in *re-6* palisade mesophyll is caused, at least in part, by a reduction in the cellular division rate early on in leaf development.

Throughout leaf development, *re-6* palisade cells appear to be bigger than those of the wild-type, although this difference only becomes significant from 14 DAS onward (Fig. 5C). By 23 DAS, *re-6* palisade cells are on average 1.8 times larger than those of the wild-type. Given that from 6 DAS onward palisade cell density was lower in *re-6* (Fig. 5B), but clear differences in cell size emerge only from 14 DAS onward (Fig. 5C), it is possible that the enlarged cell size phenotype in the *re-6* palisade arises as an indirect effect to compensate for the reduced palisade cell division rate early in leaf development or that RE has an effect on cell division early on and an effect on cell expansion later.

To further characterise how cell morphology changes over time, we quantified three shape descriptors for palisade cells of *re-6* and wild-type in the paradermal plane: aspect ratio, circularity and solidity. Aspect ratio is defined as the ratio of the major to the minor axis of each cell, such that higher values indicate more elongated cells. Circularity is calculated as (4π·area)/perimeter^2^; a value of 1 indicates a perfect circle, whereas values approaching 0 indicate increasingly elongated or irregular shapes. Solidity is defined as the ratio of the cell area to its convex area, the latter being the area of the smallest convex polygon that fully encloses the cell. A solidity of 1 indicates that a cell has no indentations, while values <1 reflect the presence of lobes, concavities, or irregular boundaries. Wild-type palisade cells start off as small, densely packed polyhedral cells, and gradually acquire their characteristic columnar shape, which looks roughly like a circle in a transverse cut (Fig. 2A). The palisade cells in *re-6* initially also follow this developmental trajectory, however, after 14 DAS their aspect ratio values increase, becoming notably more elongated than those of the wild-type (Fig. 5D). In terms of the circularity, *re-6* palisade cells were more circular than wild-type cells up until 10 DAS, probably because they are less tightly packed in *re-6*. But after 12 DAS, instead of plateauing like the wild-type cells, the circularity values of *re-6* palisade cells dropped significantly (Fig. 5E), showing that the palisade of *re-6* undergoes drastic cell morphological changes. Likewise, this trend was mirrored by the solidity values (Fig. 5F), demonstrating the emergence of deep lobes and concavities in *re-6* palisade cells. Although visually the palisade cells of *re-6* seem to acquire a spongy-like morphology at 23 DAS (Fig. 2A,E), the values for the three shape descriptors were significantly different between the palisade mesophyll cells of *re-6* and the spongy mesophyll cells of wild-type at 23 DAS (*P*=0.00065 for aspect ratio, *P*=0.00007 for circularity and *P*=0.00002 for solidity), consistent with the hypothesis that palisade mesophyll identity in *re-6* has not switched to spongy mesophyll. Together, the quantification of these shape descriptors throughout palisade development show that *re-6* palisade cells undergo progressive morphological changes that complement the observed reduction in cell density and division rate, and the increase in IAS. Thus, indicating that the *re-6* mutation not only decreases palisade cell proliferation but also disrupts palisade differentiation, leading to enlarged, irregularly shaped cells in the mutant.

To continue with the quantitative characterisation of the spongy mesophyll of *re-6*, we performed the same analysis on the first rosette leaf pair of *A. thaliana* wild-type and *re-6* plants at 23 DAS (Fig. S2), as described above for the palisade. Consistent with the qualitative observations shown in Fig. 2E and the wild-type-like patterning of spongy mesophyll porosity throughout leaf development (Fig. 2C), spongy mesophyll cell density at 23 DAS remained wild-type-like (Fig. S2A). Likewise, spongy mesophyll cell area (Fig. S2B), circularity (Fig. S2D), and solidity (Fig. S2E) remained unchanged in *re-6*. In contrast, cell aspect ratio was slightly but significantly higher in *re-6* than in the wild-type (*P*=0.0494; Fig. S2C). Together, these findings indicate that, unlike the palisade mesophyll, the spongy mesophyll is largely unaffected by the *re-6* mutation, exhibiting only a subtle change in cell shape at 23 DAS.

## DISCUSSION

Here we show that RE is required for normal palisade mesophyll morphogenesis: in *re-6*, reduced proliferation in the palisade is followed by increased cell area and the development of irregularly-shaped, lobed cells, while spongy mesophyll morphogenesis occurs as normal. These changes occur without altered stomatal density, with no detectable increase in palisade cell death, and without a spatial shift in molecular markers of cell identity. Together, our findings demonstrate that *RE* is required for palisade mesophyll morphogenesis and IAS patterning.

### RE is specifically required for normal palisade mesophyll morphogenesis

The observation that loss of RE function affects palisade, but not spongy, mesophyll morphogenesis suggests that these cell types differ in their dependence on RE-mediated plastid function. This selectivity occurs even though *RE* expression is not restricted to the palisade mesophyll (Pérez-Pérez *et al*., 2013), raising the question of how a broadly expressed gene can specifically affect morphogenesis of one cell type. There is precedent for molecular mechanisms that regulate air space patterning differentially between the spongy and palisade: our recent work showed that global alterations in brassinosteroid signalling affect IAS expansion in the palisade, but not the spongy mesophyll (Fitzsimons *et al*., 2026). Similarly, changes in stomatal density specifically alter palisade mesophyll cell density (Masle *et al*., 2005; Lee *et al*., 2015; Dow *et al*., 2017; Lehmeier *et al*., 2017; Lundgren *et al*., 2019).

In the *re-6* mutant, palisade cells acquire a more spongy-like morphology but retain *IQD22* expression and lack detectable *CORI3* expression. These expression patterns and the fact that *re-6* palisade cell shape is still significantly different to wild-type spongy mesophyll cell morphology suggest that there is no wholesale conversion of palisade cell identity and highlight that palisade cell identity and morphology can at least be partially uncoupled. Overall, these data suggest that RE may act downstream of palisade cell identity to control palisade morphogenesis, although its mode of action remains unknown.

### *RE* regulates air space patterning via palisade proliferation, independently of cell death and stomatal density

The data presented here show that RE controls early palisade proliferation: in *re-6*, cell density and tissue porosity diverge from wild-type around 6 DAS, and clonal analysis demonstrates reduced cell proliferation around this time. As increased cell size becomes significant at around 14 DAS, it is possible that RE primarily affects early palisade cell division, with later effects on cell morphology arising as indirect results of compensation for reduced cell number, as overall leaf size is maintained (Ferjani *et al*., 2007). However, it is also possible that RE affects both directly.

Previous studies in *A. thaliana* and wheat have shown that the degree of mesophyll porosity is positively correlated with the presence of open functional stomata (Dow *et al*., 2017; Lundgren *et al*., 2019). In *re-6* the density of functional stomata in the adaxial and abaxial epidermis is comparable to the wild-type, but IAS is enlarged. In addition, IAS enlargement begins before stomata are open. Therefore it is unlikely that RE affects IAS in the palisade mesophyll indirectly via a change in stomatal density.

Other studies have proposed that increased IAS volume in the *re* mutant is caused by mesophyll cell death due to increased reactive oxygen species (ROS) in mature leaves (Barth & Conklin, 2003; Overmyer *et al*., 2008; Pérez-Pérez *et al*., 2013). However, none of these studies documented any palisade cell death during *re-6* leaf development. Here, we show that, during early leaf development where air spaces diverge from wild-type, cell death is not accountable for the enlarged air spaces in the palisade of *re-6*. Instead, our findings suggest that enlarged air spaces in *re-6* are due to the reduced proliferation rate of the palisade cells early on in leaf development.

### Linking plastid amino acid homeostasis to palisade development

The recent identification of RE as a chloroplast amino acid transporter provides a biochemical starting point to understand its developmental function. RE transports arginine, lysine, ornithine and citrulline across the chloroplast inner envelope, and mutants have disrupted amino acid homeostasis (Kuhnert *et al*., 2025).

One possible connection to cell proliferation is through polyamine metabolism. In *A. thaliana*, polyamines such as putrescine and spermidine are synthesised from arginine through the arginine decarboxylase pathway (Hanfrey *et al*., 2001). Polyamines have been linked to cell proliferation in tobacco leaves (Paschalidis & Roubelakis-Angelakis, 2005), in mesophyll-derived cell cultures of several species (Tiburcio *et al*., 1986; Majewska-Sawka *et al*., 1997) and in *A. thaliana* roots (Ko *et al*., 2026). Therefore, disruption of arginine homeostasis in *re-6* could alter polyamine levels and reduce early palisade cell proliferation. Quantifying polyamine abundance and function in a cell type-specific manner during early leaf development would determine whether this pathway contributes to the *re* phenotype and if a limitation in the supply of polyamines would affect the development of the palisade but not the spongy mesophyll.

The cellular site of RE action is also currently unresolved. High levels of expression in the bundle sheath have led to the hypothesis that the bundle sheath supplies either essential metabolites (Rosar *et al*., 2012) or a differentiation signal for mesophyll development (Knappe *et al*., 2003; Pérez-Pérez *et al*., 2013; reviewed in Lundquist *et al*., 2014). Alternatively, RE may act within palisade cells themselves to control development cell-autonomously. Tissue- and clonal sector-specific rescue would help distinguish these possibilities, while single-cell metabolomics and transcriptomics of *re* may identify specific plastid metabolic defects that are associated with altered leaf structure and show how RE affects development differently in the palisade and spongy mesophyll.

In addition, since the reticulate phenotype of *re* can be rescued when grown in a short-day photoperiod (Overmyer *et al*., 2008; Pérez-Pérez *et al*., 2013) and under very low light (Barth & Conklin, 2003; Overmyer *et al*., 2008), it is possible that under these conditions the metabolic demand of the developing leaf is reduced, allowing the system to compensate and thereby suppress the reticulate phenotype. Thus, it is plausible that RE function links the regulation of palisade development in response to light levels and the metabolic state of the cell.

## CONCLUSIONS

Overall, our findings demonstrate that RE regulates palisade mesophyll morphogenesis and IAS patterning by promoting early palisade cell division and regulating later cell expansion and morphology. Understanding how RE-dependent plastid amino acid homeostasis regulates these processes will help reveal how chloroplast metabolism shapes leaf development. More broadly, future work in this area may reveal how chloroplasts integrate environmental signals to shape plastic leaf development.

## MATERIALS AND METHODS

### Plant material and growth conditions

The Col-0 wild-type and *re-6* mutant plants were obtained from the Nottingham Arabidopsis Stock Centre (NASC). To generate transgenic plants for sector analysis, wild-type plants were transformed with the 71891 plasmid by floral dip and lines selected based on resistance to BASTA and the ability to produce fluorescent sectors after heat shock. A sector line was then crossed to *re-6* to generate sectors in the *re-6* mutant background.

Prior to sowing, seeds were sterilised by incubating first for 10 min with 70% ethanol and 0.05% (w/v) sodium dodecyl sulfate (SDS) and later for 5 min with 100% ethanol. Plants were grown on plates containing MS media [0.441% Murashige and Skoog including vitamins, 0.05% (w/v) MES, 0.8% Difco agar, pH to 5.7]. Sterilised seeds were stratified in the dark at 4°C for 2 days before growing in a controlled environment chamber at 20°C, 50% relative humidity and 16 h of illumination (150 µmol·m^−2^·s^−1^).

### Construct generation

The *IQD22::GUS-mCit* and *CORI3::GUS-mCit* lines were obtained from Carl Procko (Procko *et al*., 2022) and crossed with the *re-6* mutant to generate, respectively, palisade and spongy mesophyll-specific marker lines in the *re-6* background. Heterozygous plants were imaged for *IQD22::GUS-mCit*/*re-6*, and homozygous plants were imaged for *CORI3::GUS-mCit*/*re-6*.

The heatshock sector construct 71891 was generated by golden gate cloning (see Table S1 for cloning scheme, and supplemental file 71891.gb for construct sequence). Construct design was based on the brainbow 1.0 construct where induction generates mutually exclusive sectors of either YFP or RFP (Livet *et al*. (2007), Fig. S3).

### Staining methods and confocal imaging

Nile Red dye (Sigma-Aldrich) was dissolved in 1.0 cs Silicon oil (Aston Chemicals) by shaking overnight to a final concentration of 10 µM. Leaves were stained by soaking in Nile Red solution for 10 min before mounting on a slide under a coverslip and imaging on a Leica SP8 confocal microscope using a 20× dry objective. Nile Red bound to cell membranes was excited with a 561 nm wavelength laser and visualised between 565 and 640 nm, and Nile Red free in solution was excited with a 488 nm wavelength laser and visualised between 490 and 560 nm.

Staining with propidium iodide (PI) was performed by incubating *A. thaliana* seedlings with shaking for 2 h in three parts 100% ethanol and one part glacial acetic acid, then we washed the seedlings in 10x PBS, followed by staining with 250 ng/mL of PI for 40 min and then leaving the seedlings in ClearSee (Kurihara *et al*., 2015) overnight. PI was excited with a 561 nm wavelength laser and visualised between 590 and 650 nm. To quantify leaf porosity (percentage air space in a region of tissue) unbound Nile Red fluorescence or areas with no PI fluorescence were quantified by automatic thresholding in ImageJ as described previously (Kawase *et al*., 2015, 2016), and porosity was defined as the percentage area of a 2D region that contained air space. To accurately identify palisade and spongy mesophyll tissue through-out development before defining morphological characteristics emerged, palisade and spongy mesophyll tissue were defined by position: palisade as the most adaxial layer of subepidermal mesophyll cells and spongy as the most abaxial layer of subepidermal mesophyll cells.

In leaf tissue, dead cells were visualised by staining with Sytox Orange, as described previously by Truernit & Haseloff (2008). Sytox Orange (Molecular Probes, Eugene, USA) was dissolved in DMSO and diluted to a concentration of 250 nM in water. Leaves were stained by soaking in Sytox Orange for 5 min before mounting on a slide under a coverslip and imaging on a Leica SP8 confocal microscope using a 20× dry objective. Sytox Orange was excited with a 568 nm wavelength laser and visualised between 580 and 610 nm.

To confirm that Sytox Orange could stain dead cells in the mesophyll, laser ablations were performed on the Leica SP8 using the ROI function, focusing a laser of 568 nm wavelength for 30 seconds on a single point. This caused the cell to rupture and the dye to penetrate and fluoresce. Ablation experiments with Sytox Orange were conducted at 8 DAS and 14 DAS.

To visualise *IQD22::mCit* and *CORI3::mCit* expression in mesophyll cells, leaves were embedded in 3% agarose and sectioned using a vibratome (Leica VT 1200S) to a thickness of 150 µm. Sections were stained for 10 minutes in a 1% calcofluor white solution then rinsed in water and visualised using a Leica SP8 confocal microscope. Calcofluor white was excited with a 405 nm wavelength laser and visualised between 450 and 480 nm. mCitrine was excited with a 514 nm wavelength laser and visualised between 524 and 575 nm. Chlorophyll autofluorescence was observed by exciting the sample with a 405 nm wavelength laser and visualised between 680 and 740 nm.

### Leaf area and cell counts

Whole leaves were imaged using a Keyence VHX8000 microscope, and leaf area quantified using ImageJ. Cell density measurements were calculated by using the CellposeSAM v4.0.4 cellular segmentation software (Pachitariu *et al*., 2025) to generate a ROI from confocal images of stained leaves like the ones from Fig. 2A,E. The ROI was then imported to ImageJ to obtain a cells/mm^2^ value for each leaf. Quantification of cell area and shape descriptors were performed in ImageJ using the measurements command with the ROI generated on CellposeSAM v4.0.4. Stomata density measurements were calculated by manually counting stomata number in a 500×500 µm region of epidermis tissue from a leaf stained with Nile Red. Cell death proportions in the palisade mesophyll were calculated by counting Sytox Orange stained and non-stained cells in a 500×500 µm region of the palisade mesophyll.

### Sector analysis and heat shock induction

Expression of CRE recombinase was transiently induced in 4-day old plate-grown seedlings transformed with the 71891 plasmid by heat shocking them for 60-90 seconds by floating plates in a 39°C water bath. Plants were then returned to the growth room and left to grow. Heat shocks of this length cause CRE expression in very few cells and generate clonal sectors. In individual cells that were induced, mutually exclusive clonal sectors of YFP/RFP were detected using a Leica SP8 confocal microscope and cell numbers per clone were manually counted 3 days post-induction.

### Statistical analyses and figure design

Statistical analyses, graph plotting and figure design were performed in R.

## ACKNOWLEDGEMENTS

We thank the facilities and technical staff for general support.

## AUTHOR CONTRIBUTIONS

Conceptualisation: E.A.L.B., E.A., C.D.W.; Funding acquisition: C.D.W.; Investigation: E.A.L.B., E.A., J.M.F., J.C.D., C.D.W.; Methodology: E.A.L.B., E.A., C.D.W.; Project administration: C.D.W.; Resources: E.A., J.M.F., J.C.D.; Supervision: E.A.L.B., E.A., C.D.W.; Visualisation: E.A.L.B., E.A., C.D.W.; Writing – original draft: E.A.L.B.; Writing – review & editing: E.A.L.B., E.A., J.C.D., C.D.W.

## REFERENCES

Avery Jr GS. 1933. Structure and development of the tobacco leaf. American Journal of Botany: 565–592.

Baillie AL, Fleming AJ. 2020. The developmental relationship between stomata and mesophyll airspace. New Phytologist 225: 1120–1126.

Bar M, Ori N. 2014. Leaf development and morphogenesis. Development 141: 4219–4230.

Barth C, Conklin PL. 2003. The lower cell density of leaf parenchyma in the arabidopsis thaliana mutant lcd1-1 is associated with increased sensitivity to ozone and virulent pseudomonas syringae. The Plant Journal 35: 206–218.

Chatterjee M, Sparvoli S, Edmunds C, Garosi P, Findlay K, Martin C. 1996. DAG, a gene required for chloroplast differentiation and palisade development in antirrhinum majus. The EMBO journal 15: 4194–4207.

DeLucia E, Nelson K, Vogelmann T, Smith W. 1996. Contribution of intercellular reflectance to photosynthesis in shade leaves. Plant, Cell & Environment 19: 159–170.

Donnelly PM, Bonetta D, Tsukaya H, Dengler RE, Dengler NG. 1999. Cell cycling and cell enlargement in developing leaves of arabidopsis. Developmental biology 215: 407–419.

Dow GJ, Berry JA, Bergmann DC. 2017. Disruption of stomatal lineage signaling or transcriptional regulators has differential effects on mesophyll development, but maintains coordination of gas exchange. New Phytologist 216: 69–75.

Earles JM, Théroux-Rancourt G, Roddy AB, Gilbert ME, McElrone AJ, Brodersen CR. 2018. Beyond porosity: 3D leaf intercellular airspace traits that impact mesophyll conductance. Plant Physiology 178: 148–162.

Ferjani A, Horiguchi G, Yano S, Tsukaya H. 2007. Analysis of leaf development in fugu mutants of arabidopsis reveals three compensation modes that modulate cell expansion in determinate organs. Plant Physiology 144: 988–999.

Fitzsimons JM, Rock AB, De Falbe RL, Fox S, Whitewoods CD. 2026. Brassinosteroid controls leaf air space patterning non-cell autonomously by promoting epidermal growth. Development 153: dev205110.

Fox S, Southam P, Pantin F, Kennaway R, Robinson S, Castorina G, Sánchez-Corrales YE, Sablowski R, Chan J, Grieneisen V, et al. 2018. Spatiotemporal coordination of cell division and growth during organ morphogenesis. PLoS biology 16: e2005952.

González-Bayon R, Kinsman EA, Quesada V, Vera A, Robles P, Ponce MR, Pyke KA, Micol JL. 2006. Mutations in the RETICULATA gene dramatically alter internal architecture but have little effect on overall organ shape in arabidopsis leaves. Journal of experimental botany 57: 3019–3031.

Hanfrey C, Sommer S, Mayer MJ, Burtin D, Michael AJ. 2001. Arabidopsis polyamine biosynthesis: Absence of ornithine decarboxylase and the mechanism of arginine decarboxylase activity. The Plant Journal 27: 551–560.

Hricová A, Quesada V, Micol JL. 2006. The SCABRA3 nuclear gene encodes the plastid RpoTp RNA polymerase, which is required for chloroplast biogenesis and mesophyll cell proliferation in arabidopsis. Plant physiology 141: 942–956.

Jarvis M, Briggs S, Knox J. 2003. Intercellular adhesion and cell separation in plants. Plant, Cell & Environment 26: 977–989.

Jeffree C, Dale J, Fry S. 1986. The genesis of intercellular spaces in developing leaves of phaseolus vulgaris l. Protoplasma 132: 90–98.

Kawase T, Sugano SS, Shimada T, Hara-Nishimura I. 2015. A direction-selective local-thresholding method, DSLT, in combination with a dye-based method for automated three-dimensional segmentation of cells and airspaces in developing leaves. The Plant Journal 81: 357–366.

Kawase T, Sugano SS, Shimada T, Hara-Nishimura I. 2016. Differential and simultaneous visualization of cells and airspaces in plant leaves. Bio-protocol 6: e1826–e1826.

Keddie JS, Carroll B, Jones J, Gruissem W. 1996. The DCL gene of tomato is required for chloroplast development and palisade cell morphogenesis in leaves. The EMBO journal 15: 4208–4217.

Knappe S, Löttgert T, Schneider A, Voll L, Flügge U-I, Fischer K. 2003. Characterization of two functional phosphoenolpyruvate/phosphate translocator (PPT) genes in arabidopsis– AtPPT1 may be involved in the provision of signals for correct mesophyll development. The Plant Journal 36: 411–420.

Ko D, Ruonala R, Faille A, Hellmann E, Help H, Liu H, Nielsen R, Haakonsson A, De Diego N, Paatero A, et al. 2026. Recruitment of bifunctional regulator thermospermine to methylated ribosomes directs xylem fate. Science 391: 694–699.

Kuchen EE, Fox S, Barbier de Reuille P, Kennaway R, Bensmihen S, Avondo J, Calder GM, Southam P, Robinson S, Bangham A, et al. 2012. Generation of leaf shape through early patterns of growth and tissue polarity. Science 335: 1092–1096.

Kuhnert F, Westhoff P, Valencia V, Krüger S, Vogel K, Lundquist PK, Rosar C, Goss T, Weber AP. 2025. RETICULATA1 is a plastid-localized basic amino acid transporter. Nature Plants 11: 1890–1902.

Kurihara D, Mizuta Y, Sato Y, Higashiyama T. 2015. ClearSee: A rapid optical clearing reagent for whole-plant fluorescence imaging. Development 142: 4168–4179.

Lee KJ, Bushell C, Koide Y, Fozard JA, Piao C, Yu M, Newman J, Whitewoods C, Avondo J, Kennaway R, et al. 2019. Shaping of a three-dimensional carnivorous trap through modulation of a planar growth mechanism. PLoS Biology 17: e3000427.

Lee JS, Hnilova M, Maes M, Lin Y-CL, Putarjunan A, Han S-K, Avila J, Torii KU. 2015. Competitive binding of antagonistic peptides fine-tunes stomatal patterning. Nature 522: 439–443.

Lehmeier C, Pajor R, Lundgren MR, Mathers A, Sloan J, Bauch M, Mitchell A, Bellasio C, Green A, Bouyer D, et al. 2017. Cell density and airspace patterning in the leaf can be manipulated to increase leaf photosynthetic capacity. The Plant Journal 92: 981–994.

Li H, Culligan K, Dixon RA, Chory J. 1995. CUE1: A mesophyll cell-specific positive regulator of light-controlled gene expression in arabidopsis. The Plant Cell 7: 1599–1610.

Livet J, Weissman TA, Kang H, Draft RW, Lu J, Bennis RA, Sanes JR, Lichtman JW. 2007. Transgenic strategies for combinatorial expression of fluorescent proteins in the nervous system. Nature 450: 56–62.

Lundgren MR, Mathers A, Baillie AL, Dunn J, Wilson MJ, Hunt L, Pajor R, Fradera-Soler M, Rolfe S, Osborne CP, et al. 2019. Mesophyll porosity is modulated by the presence of functional stomata. Nature Communications 10: 2825.

Lundquist PK, Rosar C, Bräutigam A, Weber AP. 2014. Plastid signals and the bundle sheath: Mesophyll development in reticulate mutants. Molecular Plant 7: 14–29.

Majewska-Sawka A, Niklas A, Jażdżewska E. 1997. The effect of polyamines on the development of sugar beet protoplasts. Biologia plantarum 39: 561–567.

Masle J, Gilmore SR, Farquhar GD. 2005. The ERECTA gene regulates plant transpiration efficiency in arabidopsis. Nature 436: 866–870.

Mathers AW, Hepworth C, Baillie AL, Sloan J, Jones H, Lundgren M, Fleming AJ, Mooney SJ, Sturrock CJ. 2018. Investigating the microstructure of plant leaves in 3D with lab-based x-ray computed tomography. Plant methods 14: 99.

Overmyer K, Kollist H, Tuominen H, Betz C, Langebartels C, Wingsle G, Kangasjärvi S, Brader G, Mullineaux P, Kangasjärvi J. 2008. Complex phenotypic profiles leading to ozone sensitivity in arabidopsis thaliana mutants. Plant, cell & environment 31: 1237–1249.

Pachitariu M, Rariden M, Stringer C. 2025. Cellpose-sam: Superhuman generalization for cellular segmentation. bioRxiv.

Paschalidis KA, Roubelakis-Angelakis KA. 2005. Sites and regulation of polyamine catabolism in the tobacco plant. Correlations with cell division/expansion, cell cycle progression, and vascular development. Plant Physiology 138: 2174–2184.

Pérez-Pérez JM, Esteve-Bruna D, González-Bayón R, Kangasjärvi S, Caldana C, Hannah MA, Willmitzer L, Ponce MR, Micol JL. 2013. Functional redundancy and divergence within the arabidopsis RETICULATA-RELATED gene family. Plant physiology 162: 589–603.

Procko C, Lee T, Borsuk A, Bargmann BO, Dabi T, Nery JR, Estelle M, Baird L, O’Connor C, Brodersen C, et al. 2022. Leaf cell-specific and single-cell transcriptional profiling reveals a role for the palisade layer in UV light protection. The Plant Cell 34: 3261–3279.

Pyke K, Marrison J, Leech A. 1991. Temporal and spatial development of the cells of the expanding first leaf of arabidopsis thaliana (l.) heynh. Journal of Experimental Botany 42: 1407–1416.

Rédei GP, Hironyo Y. 1964. Linkage studies. Arabidopsis Information Service 1: 9–10.

Rosar C, Kanonenberg K, Nanda AM, Mielewczik M, Bräutigam A, Novák O, Strnad M, Walter A, Weber AP. 2012. The leaf reticulate mutant dov1 is impaired in the first step of purine metabolism. Molecular plant 5: 1227–1241.

Schmidt R, Kunkowska AB, Schippers JH. 2016. Role of reactive oxygen species during cell expansion in leaves. Plant Physiology 172: 2098–2106.

Slaton MR, Smith WK. 2002. Mesophyll architecture and cell exposure to intercellular air space in alpine, desert, and forest species. International Journal of Plant Sciences 163: 937–948.

Smith WK, Vogelmann TC, DeLucia EH, Bell DT, Shepherd KA. 1997. Leaf form and photosynthesis. BioScience 47: 785–793.

Terashima I, Inoue Y. 1984. Comparative photosynthetic properties of palisade tissue chloroplasts and spongy tissue chloroplasts of camellia japonica l.: Functional adjustment of the photosynthetic apparatus to light environment within a leaf. Plant and Cell Physiology 25: 555–563.

Tiburcio AF, Kaur-Sawhney R, Galston AW. 1986. Polyamine metabolism and osmotic stress: II. Improvement of oat protoplasts by an inhibitor of arginine decarboxylase. Plant Physiology 82: 375–378.

Truernit E, Haseloff J. 2008. A simple way to identify non-viable cells within living plant tissue using confocal microscopy. Plant methods 4: 15.

Van Lijsebettens M, Clarke J. 1998. Leaf development in arabidopsis. Plant Physiology and Biochemistry 36: 47–60.

Voll L, Häusler RE, Hecker R, Weber A, Weissenböck G, Fiene G, Waffenschmidt S, Flügge U-I. 2003. The phenotype of the arabidopsis cue1 mutant is not simply caused by a general restriction of the shikimate pathway. The Plant Journal 36: 301–317.

Zhang L, McEvoy D, Le Y, Ambrose C. 2021. Live imaging of microtubule organization, cell expansion, and intercellular space formation in arabidopsis leaf spongy mesophyll cells. The Plant Cell 33: 623–641.

