## Supplementary data for "The chloroplast-localised transporter RETICULATA controls palisade mesophyll cell division and expansion"

### Supplementary information

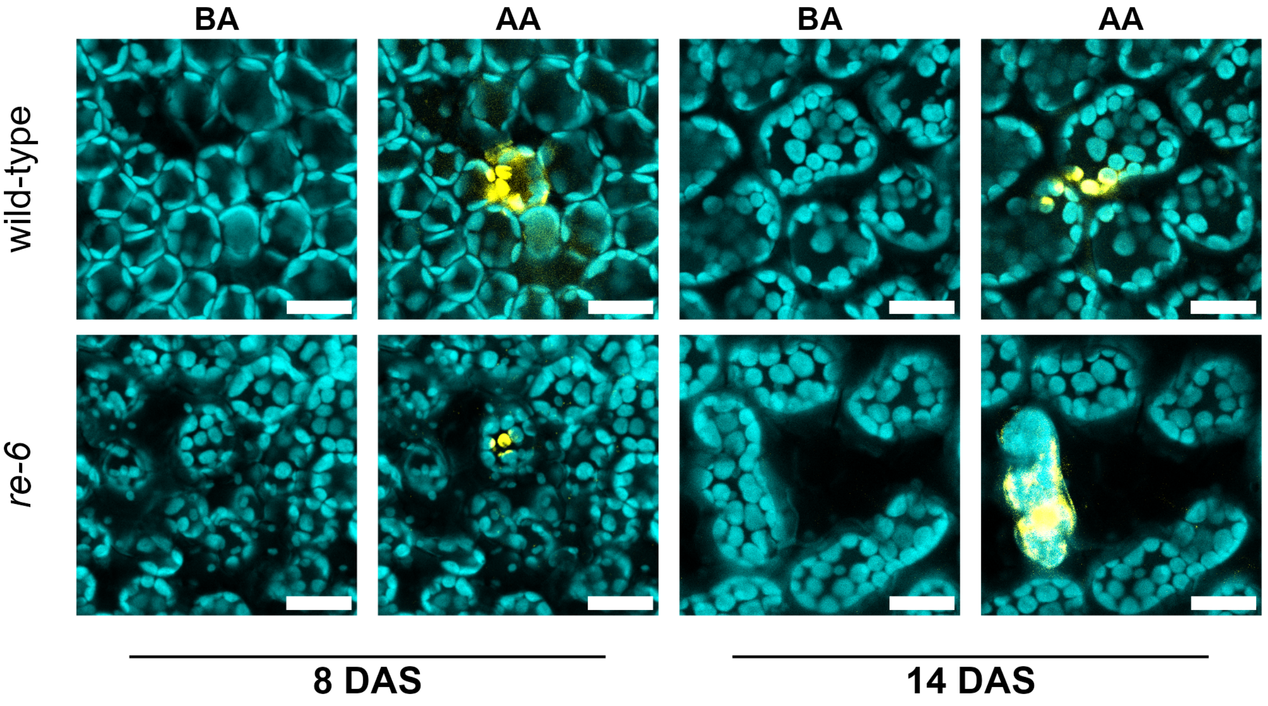


**Fig. S1. Sytox Orange stains palisade mesophyll cells after targeted laser ablation.** Representative images before (BA) and after ablation (AA) of palisade mesophyll of rosette leaf one stained with Sytox Orange at 8 and 14 DAS. wild-type (top row) and *re-6* (bottom row). Chloroplast autofluorescence is visualised in cyan and Sytox Orange in yellow. Scale bars: 20 μm.

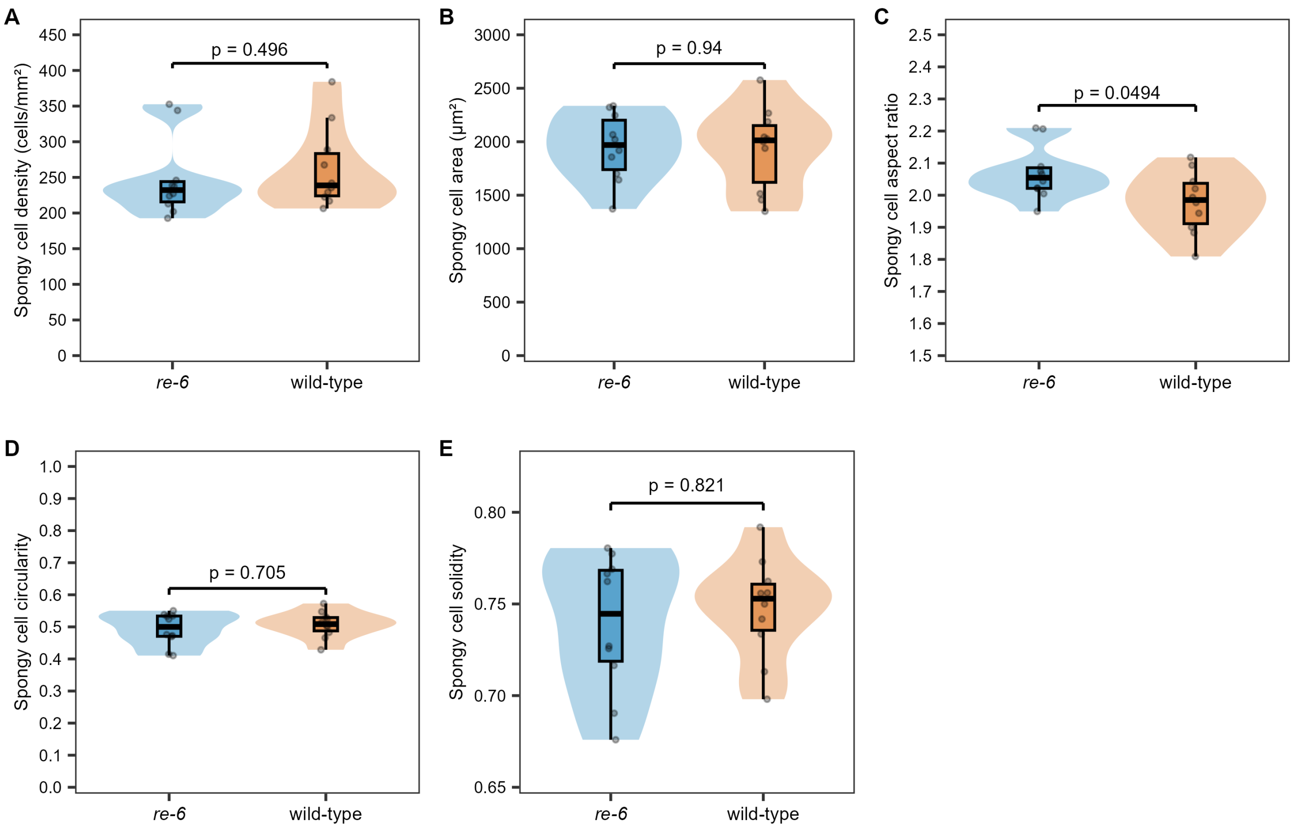


**Fig. S2. The size, morphology and division rate of *re-6* spongy mesophyll cells is wild-type-like.** (A–F) Spongy cell (A) density (cells/mm$^2$), (B) area (μm$^2$), (C) aspect ratio, (D) circularity and (E) solidity of wild-type and *re-6* first rosette leaf pair at 23 DAS. Each data point represents the mean value for a leaf. Statistical significance was calculated with the Kruskal-Wallis test.

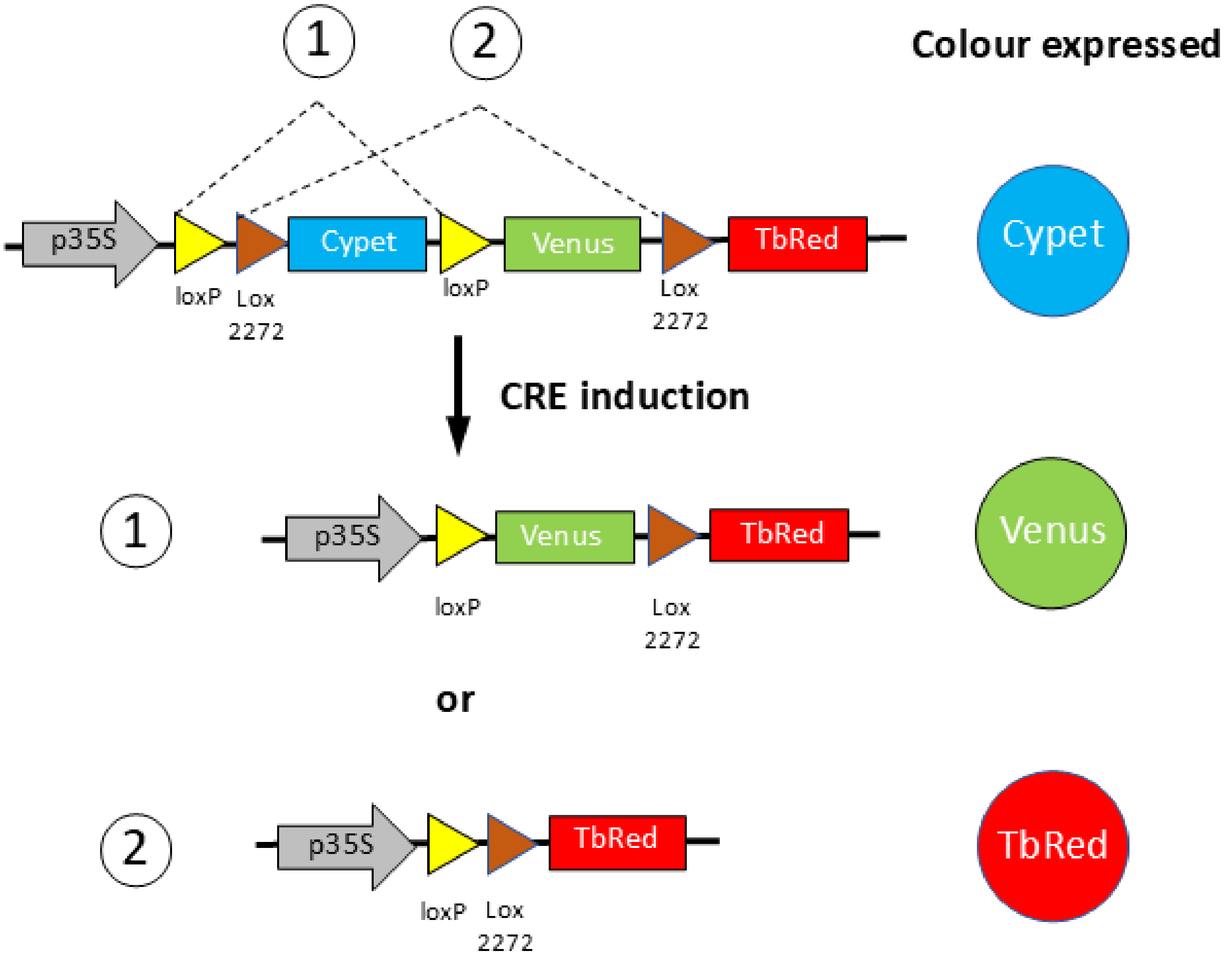


**Fig. S3. Diagram of the heat-shock inducible CRE-lox recombinase system used to study cell division rates in the palisade mesophyll.** Heat-shock induced CRE-mediated recombination of lox sites generates mutually exclusive sectors expressing Venus and TbRed in plants transformed with construct 71891. Heat-shock induces expression of CRE recombinase, which causes recombination between either loxP sites to cause Venus expression (1), or lox2272 sites to drive TurboRed expression (2).
